# Antimicrobial Susceptibility Profile of *Neisseria gonorrhoeae* from Patients Attending a Medical Laboratory, Institut Pasteur de Madagascar between 2014-2020: Phenotypic and Genomic characterization in a Subset of *Neisseria gonorrhoeae* Isolates

**DOI:** 10.1101/2023.04.20.537661

**Authors:** Lala Fanomezantsoa Rafetrarivony, Mamitina Alain Noah Rabenandrasana, Elisoa Ratsima Hariniaina, Frédérique Randrianirina, Anthony Marius Smith, Tania Crucitti

## Abstract

2

The antimicrobial resistance of *Neisseria gonorrhoea* to all classes of current available antibiotics is a global concern. National surveillance programmes monitoring the susceptibility profiles of *Neisseria gonorrhoeae* hardly exist in resource constraint settings. Therefore, little is known about the antimicrobial susceptibility profile and associated genetic resistance mechanisms of *N. gonorrhoeae* in Madagascar. We report susceptibility data of *N. gonorrhoeae* isolates obtained by the medical laboratory of the Institut Pasteur de Madagascar, from 2014 -2020. In addition, we present data on the antimicrobial resistance mechanisms and antimicrobial resistance profile of a subset of isolates (N=46), including all isolates available of 2020. Over the study period, ceftriaxone resistant isolates exceeding the threshold of 5% in 2017 and 2020, were reported. Of the subset of re-tested isolates, all were found susceptible to ceftriaxone, azithromycin, and spectinomycin. Conversely, all isolates were resistant to ciprofloxacin and the majority was also resistant to penicillin and tetracycline. None of the isolates carried the mosaic *penA* gene and chromosomal mutations associated to the antibiotic resistance in *gyrA, parC, penA, ponA, porB* and *mtrR* genes were detected. The high rate of resistance to Penicillin and Tetracycline is explained by the presence of β-lactamase *bla*_TEM_ and *tetM* genes, respectively, a plasmid mediated resistance. We found a high number of circulating multilocus sequence types. Almost half of them were new types, and one of them was among the four most predominant sequence types. Our report provides a detailed dataset obtained through phenotypic and genotypic methods which will serve as baseline for future surveillance of *N. gonorrhoeae* in our setting, and Madagascar.

**Repositories:** This Whole Genome Shotgun project has been deposited at DDBJ/ENA/GenBank under the Accession BioProject PRJNA929018.

**Impact statement:** *Neisseria gonorrhoeae* is becoming increasingly resistant to all classes of antibiotics available for infections treatment.

Resource constraint settings encounter difficulties in implementing surveillance of the antimicrobial susceptibility of *N. gonorrhoeae*. We report here antimicrobial susceptibility results from gonorrhoea among patients consulting a medical laboratory in Antananarivo, Madagascar in 2014–2020. We used whole-genome sequencing to identify resistance mechanisms in a subset of isolates including all viable isolates of 2020.

We report multilocus sequence types and discuss phenotypes and genotypes according to the phylogenetic analysis of the isolates. The susceptibility results are the first in a decade to be reported. We set the baseline to study further the evolution and transmission of *N. gonorrhoeae* resistance mechanisms and genotypes in general. Our report will enable improving surveillance of *Neisseria gonorrhoeae* in Madagascar and Africa.

Overall, it will contribute to the global, regional, and national surveillance of *N. gonorrhoeae*. In addition, it may set a benchmark for implementation in other settings facing barriers implementing phenotypic resistance surveillance of *N. gonorrhoeae*.

**Data summary:** The source code of the Tormes pipeline used in this study is also available on Github: https://github.com/nmquijada/tormes.

The SRA sequences have been deposited in the NCBI SRA database under accession numbers SRR23260223 and SRR23260193.

The genome assemblies can be accessed using the accession numbers: SAMN32949360 - SAMN32949405.

The MLST genes of all isolates can be accessed through this link: https://doi.org/10.5281/zenodo.7581537

The Perl scripts used for a quality filtering of the assemblies which is to remove small contigs can be accessed through the following link: https://github.com/Alainnoah/Remove-Small-Contigs-Draft-Assemblies.

Genomic analysis with metadata in Pathogenwatch could be accessed with this link: https://pathogen.watch/collection/f7t4wjtjlybh-mdgwhofa

A visualization of genomic epidemiology of our isolates with Microreact can be accessed with this link: https://microreact.org/project/7FADQYwhm5h1zJyJTynM6R-unnamed-project

A supplementary material in an Excel sheet summarizing the lists of plasmids used for alignment (Table S1), the statistics of the sequences (Table S2), and the minimum inhibitory concentration values of *N. gonorrhoeae* isolates and their antimicrobial resistance genetic mechanisms (Table S3) is available through this link: https://doi.org/10.6084/m9.figshare.21973511

## 5. Introduction

Bacteria are becoming increasingly resistant to antimicrobials, globally. *Neisseria gonorrhoeae*, the agent of gonorrhoea, is no exception. The bacterium developed or acquired since the use of antibiotics for treatment, mechanisms to escape their actions. Today, antibiotic-resistant *N. gonorrhoeae* is considered a menace at a global level (1, 2), and features on the research priority list of WHO and urgent threats lists of the CDC (1, 3).

According to the WHO estimates for 2016, the African region had the highest prevalence of *N. gonorrhoeae* in the world. They estimated that in 2016 gonorrhoea contributed to almost 12 million and 10 million new cases among men and women 15 to 49 years of age in Sub-Saharan Africa, respectively (4).

Untreated gonorrhoea may result in serious complications such as epididymitis in men, pelvic inflammatory disease and endometritis in women, ectopic pregnancy in pregnant women, and neonatal conjunctivitis in neonates (5, 6). Treatment is hindered because most cases are asymptomatic (7, 8) and not detected but also, more importantly, by the increasing occurrence of antimicrobial resistance of the gonococcus to all classes of antibiotics.

The antibiotic resistance mechanisms of *N. gonorrhoeae* are chromosomally mediated, acquired through mutations or horizontal gene transfer and homologous recombination, or are introduced by mobile genetic elements such as plasmids. To date, several chromosomal-mediated determinants leading to resistance were identified. The mutations in genes such as *penA*, *ponA*, *mtrR*, *porB*, *rpsJ*, *gyrA*, and *parC*, among others, may result in: alterations or modifications of the antimicrobial’s target; an increased antimicrobial influx; or a decreased antimicrobial efflux. *N. gonorrhoeae* can also obtain genes through the acquisition of plasmids. Several types of β-lactamases, hydrolysing the β-lactam ring of β-lactam antimicrobials, and β-lactamase encoding plasmids have been identified and described. *tetM* is another plasmid-born gene mediating tetracycline resistance by replacing tetracycline from its ribosome target resulting in further protein synthesis. The gene is carried by smaller conjugative plasmids. Plasmid-mediated resistance results in a sudden high level of resistance, while chromosomally-mediated resistance is acquired gradually through mutations or recombination in the chromosome (9–11). Surveillance of the antimicrobial susceptibility and circulating *N. gonorrhoeae* isolates provides data on which antibiotics to use for treatment. In addition, antibiotics showing a resistance rate below 5% can be recommended for national treatment guidelines (2).

Whole genome sequencing (WGS) has been used to identify genetic determinants and predict antimicrobial resistance. In addition, molecular typing schemes based on WGS can be used to determine the spread of the different genotypes, and describe their relatedness and evolution. Finally, WGS can guide the treatment guidelines in pointing out which antibiotics to preferably exclude from the guidelines.

Data on the antimicrobial susceptibility profiles of *N. gonorrhoeae* isolated in Madagascar is scarce. We found only three scientific papers indexed in Pubmed and published between 2002 and 2009 reporting on the antimicrobial susceptibility of *N. gonorrhoeae* in Madagascar. The latest paper presented data from 47 *N. gonorrhoeae* isolates collected from 2004 to 2006 and for which the minimal inhibitory concentrations (MICs) for penicillin (PEN) and ciprofloxacin (CIP) were determined using the E test (12). A second paper, published in 2008, reported a total of 126 isolates obtained during the same period, 2004-2006. MICs were determined by agar dilution for PEN, tetracycline (TET), CIP, ceftriaxone (CRO), and spectinomycin (SPT) (13). The oldest paper dated 2002 and presented MIC results of 46 *N. gonorrhoeae* isolated from women presenting genital discharge and 21 isolates obtained from symptomatic men. MICs were measured for PEN, TET, CIP, CRO, and SPT (14).

We aimed to describe the antimicrobial susceptibility profiles of *N. gonorrhoeae* isolated from a patient population consulting the medical laboratory (CBC) of the Institut Pasteur de Madagascar (IPM) during 2014-2020. In addition, we performed the phenotypic and genomic characterization of antimicrobial resistance in a subset of isolates including all viable isolates of 2020 to get baseline data and finally, we described briefly the molecular epidemiology of these isolates.

## 6. Methods

### Study method

From 2014 to 2020, we retrieved retrospective patients’ data collected by the CBC of the IPM, Antananarivo, Madagascar, as part of routine management. We exported coded patient and specimen data together with the MICs values of CRO, azithromycin (AZT), and CIP, as well as the β-lactamase results of the *N. gonorrhoeae* isolates from the Adagio^TM^ automated reading system.

The CBC used selective (vancomycin, colistin, amphotericin, trimethoprim) GC agar (HiMedia Laboratories, Mumbai, India) supplemented with 1% vitox (Oxoid, Hampshire, United Kingdom) and 1% haemoglobin (Oxoid, Hampshire, United Kingdom) for growth and isolation of *N. gonorrhoeae*. After incubation at 37±2°C in a 5% CO2 atmosphere for 18-48h, colonies morphologically suspected to be *N. gonorrhoeae* were identified by Matrix-assisted laser desorption/ionization time-of-flight mass spectrometry (MALDI-TOF MS). MICs for CRO, CIP and AZT were determined using E test strips (bioMérieux, Marcy-l’Etoile, France). Production of β-lactamase was detected using a cefinase disk (bioMérieux, Marcy-l’Etoile, France). All results were entered into the Adagio™ system. Isolates were stored in skim milk +20%glycerol at -80°C at the medical laboratory.

We revived and re-tested a sub-set of isolates on GC agar (HiMedia Laboratories, Mumbai, India) supplemented with 1% vitox (Oxoid, Hampshire, United Kingdom) and 1% haemoglobin (Oxoid, Hampshire, United Kingdom), incubated at 37±2°C in a candle extinction jar for 18-48h. Presumptive *N. gonorrhoeae* colonies were identified using MALDI-TOF-MS. MICs for CRO, CIP, PEN, TET, SPT, and AZT were obtained using E-test (bioMérieux, Marcy-l’Etoile, France). WHO reference gonococcal strains were used as quality control for the E-testing method.

All MIC results were interpreted according to the clinical breakpoints recommended by the Comité de l’antibiogramme de la Société Française de Microbiologie (CASFM), 2021, v1.0 April, based on the breakpoints set by the European Committee on Antimicrobial Susceptibility Testing (EUCAST).

### Whole genome sequencing

Genomic DNA was extracted from pure cultures using QIAamp DNA Mini Kit on a Qiacube automate (Qiagen GmbH, Hilden, Germany) and according to the manufacturer’s instructions. DNA was quantified using nanodrop. All DNA extracts were shipped on dry ice to the National Institute for Communicable Diseases (NICD), Johannesburg, South Africa, for whole genome sequencing on the Illumina MiSeq sequencing platform (Illumina, Inc., San Diego, CA, United States) and according to the procedures in place.

### Bioinformatic analysis

We used TORMES pipeline v1.2.1 (15) to conduct the whole genome analysis as briefly described hereafter. Prinseq and Trimmomatic were used for quality control and filtering of the reads, respectively. De novo assembly was performed using SPAdes (16). We performed assembly statistics using QUAST (17). We extracted rRNA genes from the assembly using Barrnap (T. Seemann) and classified them taxonomically using the RDP Classifier (18). Kraken2 (19) was used to infer the taxonomy of each isolate. Gene prediction and annotation were performed using Prodigal (20) and Prokka (21). We screened for antimicrobial resistance genes using ABRicate (T. Seemann) with the databases ResFinder (22), CARD (23) and ARG-ANNOT (24).

Contigs sizes less than 500bp were removed with a local script. We used the quality thresholds of assemblies as previously described (25,26).

Multi-Locus Sequence Typing (MLST) and genetic Antimicrobial Resistance (AMR) determinants were obtained from Pathogenwatch (https://pathogen.watch/).

The plasmids associated with *tetM* and *bla_TEM_* genes were identified using the Blastn program and a custom plasmid database (Supplementary Table S1). We considered a coverage and identity score of > 99% and ≥99%, respectively.

Phylogenetic analysis based on SNP alignment was performed on the Pathogenwatch platform combined with Microreact (https://microreact.org/) for visualization. We completed our study dataset with the reference sequences from WHO.

### Statistical method

Coded patient and bacteriological data comprised in the exported database were summarised using medians and interquartile ranges for continuous variables and frequencies and proportions for categorical data.

The MIC_90_ was determined as the minimal concentration of the antimicrobial required to inhibit the growth of 90% of the isolates.

## 7. Results

### Neisseria gonorrhoeae isolates

Over the study period, the CBC identified a total of 395 *N. gonorrhoeae* isolates. More than half of them (58%) were obtained from women. Overall, the age of the patients ranged from less than 1 year to 73 years, with women being slightly younger, Table 1. A total of 18 (4.6%) patients were younger than 18 years: 15 of them were girls and 3 were boys. Nine girls were below the age of 12 years, six girls were between 12 and 18 years, one boy was below 1 year and the other two boys were 14 and 17 years, respectively.

**Table 1.**
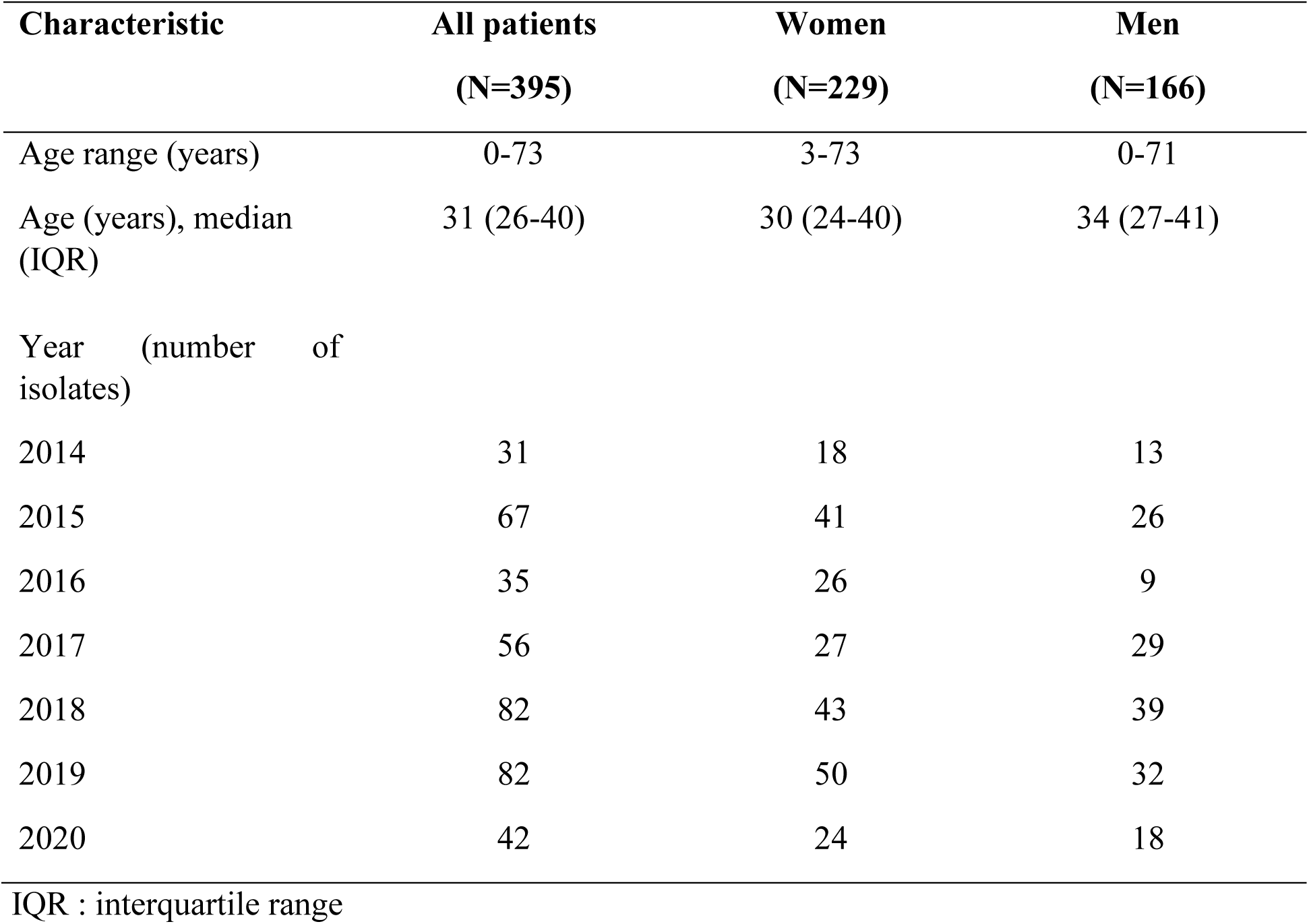
Characteristics of patients with gonorrhoea

| Characteristic | All patients<br>(N=395) | Women<br>(N=229) | Men<br>(N=166) |
| --- | --- | --- | --- |
| Age range (years) | 0-73 | 3-73 | 0-71 |
| Age (years), median (IQR) | 31 (26-40) | 30 (24-40) | 34 (27-41) |
| Year (number of isolates) |  |  |  |
| 2014 | 31 | 18 | 13 |
| 2015 | 67 | 41 | 26 |
| 2016 | 35 | 26 | 9 |
| 2017 | 56 | 27 | 29 |
| 2018 | 82 | 43 | 39 |
| 2019 | 82 | 50 | 32 |
| 2020 | 42 | 24 | 18 |

IQR : interquartile range

Almost all isolates were from genital secretions (cervicovaginal secretions in women and urethral secretions in men), followed by sperm (N=3), wound (N=1), puncture fluid (N=2), blood (N=1), and pus (N=1).

From the total amount of specimens provided for *N. gonorrhoeae* culture, the gonococcus was isolated in 4-10% of specimens from men and in 0.4-0.9% of specimens from women.

### Antimicrobial susceptibility of *Neisseria gonorrhoeae* over time

The MIC for CRO was determined in 389 isolates: 372 (96%) isolates were susceptible and 17 (4%) showed a reduced susceptibility. The resistance rate exceeded the threshold range of 5% in 2017 and 2020. Over the years the MIC_90_ varied from 0.004 to 0.02 µg/mL, with a lower concentration in the last three years of the study period, Table 2.

**Table 2.** Susceptibility to ceftriaxone and ciprofloxacin of t Neisseria gonorrhoeae over the study period

| Ceftriaxone N=372 |  |  |  |  | Ciprofloxacin N=363 |  |  |  |
| --- | --- | --- | --- | --- | --- | --- | --- | --- |
| Year | R | I | S | MIC<sub>90</sub><br>(µg/mL) | R | I | S | MIC<sub>90</sub><br>(µg/mL) |
| 2014 | 0 | 0 | 28 | 0.02 | 29 | 0 | 0 | 4 |
| 2015 | 0 | 3 | 64 | 0.006 | 46 | 0 | 0 | 3 |
| 2016 | 0 | 1 | 34 | 0.02 | 31 | 0 | 0 | 2 |
| 2017 | 0 | 4 | 50 | 0.016 | 50 | 2 | 3 | 3 |
| 2018 | 0 | 3 | 78 | 0.004 | 79 | 0 | 1 | 3 |
| 2019 | 0 | 3 | 79 | 0.006 | 81 | 0 | 0 | 4 |
| 2020 | 0 | 3 | 39 | 0.004 | 42 | 0 | 0 | 3 |

MIC: Minimum inhibitory Concentration; R: Resistant; I: Reduced susceptible; S: Susceptible

During the study period, the MIC results for CIP of a total of 364 isolates were determined, including all isolates obtained in 2020. Of the 31 isolates which were not tested, 22 of them were isolated in 2015 and counted as almost one-third of the isolates obtained that year. All isolates were resistant to ciprofloxacin except three and one susceptible isolate detected in 2017 and 2018, respectively. Two isolates obtained in 2017 showed a reduced susceptibility. The MIC_90_ values remained stable over the years (range 2-4 µg/mL).

Azithromycin was tested only in 2019 and 2020 for 95 isolates: 53 isolates in 2019 and 42 isolates in 2020. Three isolates had a MIC value > 0.5 µg/mL and were determined resistant according to CASFM, 2021. Two were detected in 2020 with a MIC of 0.64 and 0.75 µg/mL, respectively. The resistant strain obtained in 2019 had a MIC of 1.5 µg/mL against AZT. The MIC_90_ evolved from 0.064 µg/mL in 2019 to 0.32 µg/mL in 2020.

Almost all isolates produced β-lactamase, according to the nitrocefin test. Exceptions were found in 2016, 2017, and 2019 with 6%, 2% and 4% of the *N. gonorrhoeae* isolates not producing β-lactamase, respectively.

### Subset of Neisseria gonorrhoeae isolates

We retrieved from the CBC storage collection all isolates stored in 2020 and those found CRO and/or AZT resistant or reduced susceptible of previous years. We recovered 68 isolates, of them 46 were identified as *N. gonorrhoeae*. Eight stored isolates did not grow and 14 were identified as not being *N. gonorrhoeae*. Finally, the 46 *N. gonorrhoeae* isolates comprised 33/42 (71.7%) isolates from 2020, six determined reduced susceptible to CRO and two resistant to AZT.

We re-tested and determined the MICs of CRO, AZT, SPT, CIP, PEN, and TET for all 46 isolates, Table 3.

**Table 3.** Susceptibility profiles obtained after re-testing of the isolates submitted to whole genome sequencing (N=46)

| Antibiotic | 2020 (n=33) |  |  | 2014-2019 (n=13) |  |  | Breakpoints (R) EUCAST 2021 |
| --- | --- | --- | --- | --- | --- | --- | --- |
|  | R n(%) | I n(%) | S n(%) | R n(%) | I n(%) | S n(%) |  |
| **CRO** | - | - | 33 (100%) | - | - | 13 (100%) | >0,125 µg/mL |
| **AZT** | - | - | 33 (100%) | - | - | 13 (100%) | >0,5 µg/mL |
| **SPT** | - | - | 33 (100%) | - | - | 13 (100%) | >64 µg/mL |
| **CIP** | 33 (100%) | - | - | 13 (100%) | - | - | >0,06 µg/mL |
| **TET** | 27 (82%) | - | 6 (18%) | 13 (100%) | - | - | >1 µg/mL |
| **PEN** | 27 (82%) | 6 (18%) | - | 11 (85%) | 2 (15%) | - | >1 µg/mL |
| **MDR** | 23 (70%) |  |  | 11 (85%) |  |  |  |

R: Resistant; I: Reduced susceptible; S: susceptible; EUCAST: The European Committee on Antimicrobial Susceptibility Testing; CRO: Ceftriaxone; AZT: Azithromycin; SPT: Spectinomycin; CIP: Ciprofloxacin; TET: Tetracycline; PEN: Penicillin G; MDR: Multi-drug resistant (Resistant to ≥3 antibiotics)

From all isolates, the complete genome sequence was achieved with a total assembly length between 2.0 to 2.1 Mbp. (Supplementary Table S2).

None of the isolates were determined resistant or reduced susceptible to CRO using E-test. The *penA* alleles coding for the penicillin binding protein 2 were non-mosaic, the most prevalent was *penA*.14 (n=20) followed by *penA*.19 (n=12), *pen*A.2 (n=7), *penA.*22 (n=6) and *penA*.9 (n=1).

All isolates were susceptible to AZT in vitro. A 57 Adenosine deletion (57Adel) in *mtrR* promoter region was found in three isolates with a slight increase of MIC_AZT_ (0.064-0.125 µg/mL), two were combined with a mutation G45D in *mtrR* coding region. A disrupted *mtrR* was detected in one isolate with MIC_AZT_ =0.064µg/mL.

All 46 isolates were found phenotypically resistant to CIP. The accumulation of mutations in *gyrA* and *parC* contributed to the high level of MIC_CIP_ as shown in Table 4. Conversely, two isolates did not possess any mutation in the *parC* genes and displayed a low-level resistance to CIP (MIC≤0.75µg/mL).

**Table 4.** Genomic resistance mechanisms and minimal inhibitory concentration against ciprofloxacin

| *parC* | *gyrA* |  | MIC<sub>CIP</sub> (µg/mL) |
| --- | --- | --- | --- |
|  | S91F, D95A | S91F, D95G | Median (min;max) |
| WT | 1 | 1 | 0.125 / 0.75 |
| D86N | 2 | - | 1.12 (0.75; 1.5) |
| S87N, E91K | 3 | - | 3 (1.5; 4) |
| S87R | 6 | - | 1.5 (1 ; 4) |
| S87N | 33 | - | 2 (0.75; 4) |
| Total | 45 | 1 |  |

WT: Wild type; min: minimum; max: maximum

Eight (17%) isolates were reduced susceptible to PEN, and the remaining 38 (83%) were resistant, Table 3. The insertion D345 in *penA* was observed in all isolates. The SNP L421P in the *ponA* gene and the mutations in the *mtrR* coding region A39T or G45D plus 57Adel in the *mtrR* promoter, combined (n=6) or not (only L421P n=2), were carried by all isolates with reduced susceptibility to penicillin. In addition to these mutations, the SNP A121S in *porB1b* was detected in two resistant isolates. From the 38 resistant isolates, 36 contained a β-lactamase *bla*_TEM_ gene, TEM-1 in 27 isolates, TEM-135 in six isolates, and TEM-206 in three isolates, Table 5. All *bla*_TEM-135_ were carried by the Asian plasmid type.

**Table 5.** β-lactamase genes and plasmid type detected among the penicillin resistant isolates

| *bla*<sub>TEM</sub><br>allele | B-lactamase plasmid type |  |  |  | Total |
| --- | --- | --- | --- | --- | --- |
|  | African<br>type<br>(pJD5)<br>n(%) | Johannesburg<br>type (pEM1)<br>n (%) | Asian<br>type<br>(pJD4)<br>n (%) | Not<br>determined<br>n (%) | n<br>Median MIC<sub>PEN</sub> (μg/mL)<br>(min; max) |
| TEM-1 | 17<br>(59%) | 8 (27%) | - | 4 (14%) | 29<br>32 (0.125; 256) |
| TEM-135 | - | - | 6<br>(100%) | - | 6<br>72 (16 ; 256) |
| TEM-206 | 2 (67%) | - | 1 (33%) | - | 3<br>3 (2 ; 12) |
| Total | 19<br>(50%) | 8 (21%) | 7 (18%) | 4 (11%) | 38 |

PEN: Benzylpenicillin

A total of 87% (40/46) of the isolates were phenotypically resistant to TET. All isolates, except one resistant, carried a mutated *rpsJ* gene V57M. The isolate lacking the mutation, carried the *tetM* gene and an *mtrR* coding sequence with the A39T mutation. Almost all of the TET resistant isolates (39/40) contained plasmids with a *tetM* gene, and one susceptible isolate carried a disrupted *tetM* gene. The American plasmid type was the most predominant (n=30, 77%). The Dutch plasmid was identified in the other nine (23%) isolates. The only resistant isolate (MIC_TET_=12µg/mL) lacking the *tetM* gene had mutations G45D in the *mtrR* coding sequence and 57Adel in the *mtrR* promoter region.

A total of 33 isolates carried both the β-lactamase and the conjugative plasmid.

All isolates were susceptible to SPT, and all 16S rRNA genes were wild type (WT).

We did not test *in vitro* the susceptibility to sulphonamides, but *in silico* we detected a high proportion (98%) of isolates carrying the *folP* R228S mutation.

Supplementary Table S3 summarises the values of the MICs of the tested antimicrobials and associated antimicrobial resistance mechanisms of the *N. gonorrhoeae* isolates.

### Antimicrobial resistance prediction by Pathogenwatch

The MIC results were in complete accordance with the antimicrobial resistance prediction for AZT, SPT, CRO and CIP according to Pathogenwatch. Pathogenwatch predicted intermediate or reduced susceptibility for PEN in two isolates with a resistant phenotype, and predicted resistance in two other isolates with reduced susceptibility phenotype. All isolates which were defined TET resistant according to their MIC_TET_ values, were also predicted resistant by Pathogenwatch. However, of the six phenotype-susceptible isolates, five were predicted intermediate and one resistant.

### Molecular typing

Among the 46 isolates, we could identify a total of 23 MLST Sequence types (STs), of which 10 were new ST profiles. The ST9903 and a new ST with a profile (*abcZ*59, *adk*39, *aroE*67, *fumC*1167, *gdh*149, *pdhC*153, *pgm*65) called ST Unk_1, were the most prevalent STs, each of which clustered six isolates. ST1931 and 7822 included five isolates, each. ST1588 clustered three isolates and ST7827 / ST10935/ a novel ST Unk_9 (*abcZ*59, *adk*39, *aroE*67, *fumC*∼190, *gdh*148, *pdhC*153, *pgm*65) clustered each two isolates. The remaining 15 STs were singletons.

All the new STs plus eight previously defined STs were grouped into two *penA* alleles types, namely *penA*-14 and 19. ST7822 and ST9903 were associated with *penA*-2 and *penA*-22, respectively. The highly PEN resistant isolates carrying *bla*_TEM-135_ belonged exclusively to these two STs.

The MIC_AZT_ were slightly increased in ST7822 and 9903 (median= 0.047 and 0.0395µg/mL, respectively) compared to the other STs (median=0.023 µg/mL). The median of MIC_PEN_ was elevated in ST7822 and 1931 (32µg/mL) while the median of MIC_TET_ was elevated in ST1588 (median MIC_TET_=16µg/mL). ST Unk_1 showed the most elevated median of MIC_CIP_=3µg/mL. Although all isolates were susceptible to SPT, the median of MIC was slightly higher for isolates belonging to ST7827 (MIC_SPT_=8µg/mL).

The most prevalent new ST Unk-1 was also detected among isolates of 2015, and ST1931 was identified in isolates collected in 2014. A total of 18 STs was identified among the 33 isolates of 2020. Half of the isolates pertained to four STs: ST7822 (n=5); ST9903 (n=4); STUnk_1 (n=4); ST1931 (n=3). Seven STs were novel.

### Phylogenetic analysis

Genomic core analysis of the isolates combined with 14 reference gonococcal genomes (WHO, n=14) and *N. gonorrhoeae FA1090* using Pathogenwatch resulted into four groups divided into two main lineages (Figure 1).

**Figure 1.**
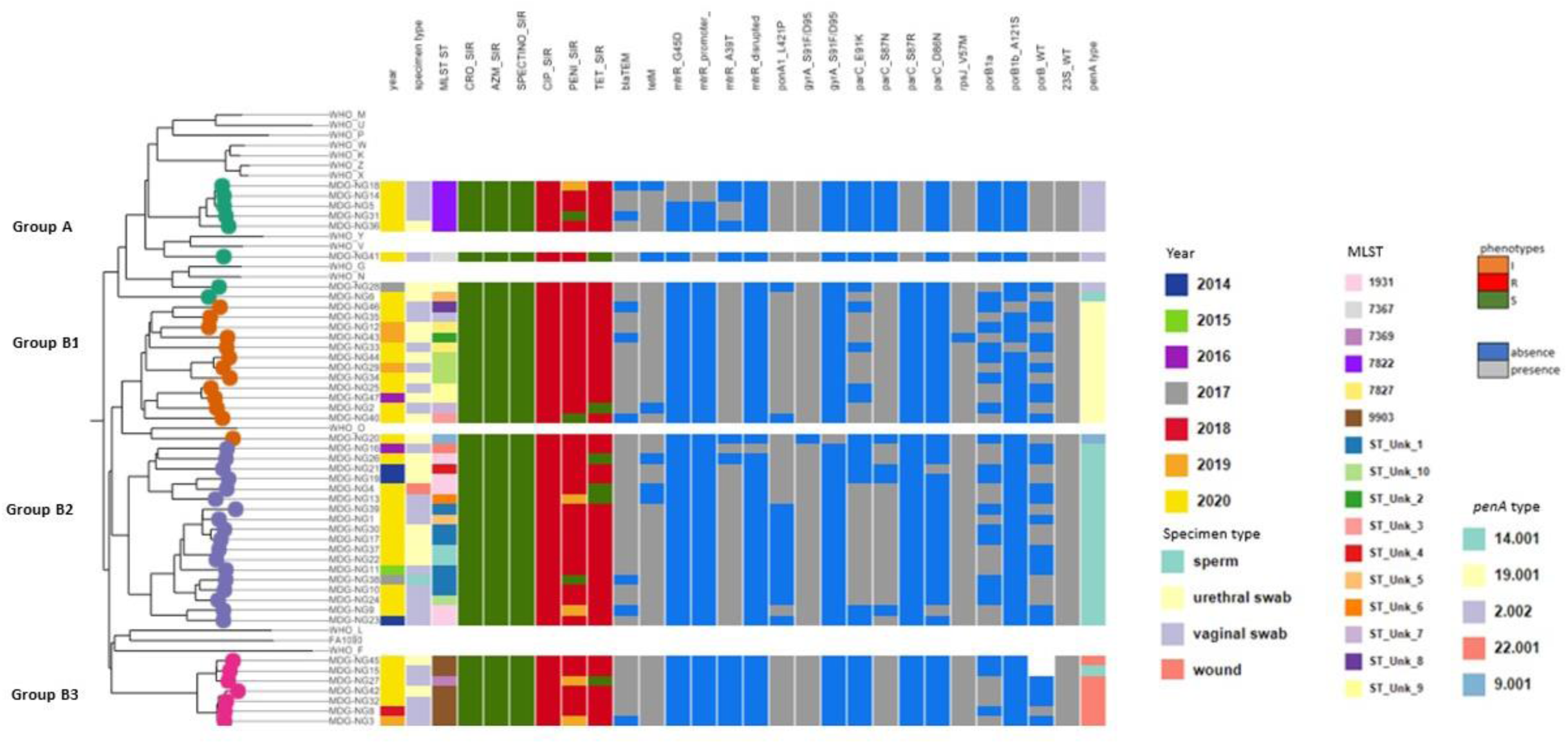
Phylogenetic tree using core genome SNP analysis for 46 GC isolates from Madagascar, 14 WHO strains, and *N.gonorrhoeae FA 1090* genome

The first lineage clustered nine isolates into one group A, while the second was clustered in three groups with 12, 18 and seven isolates corresponding to groups B1, B2 and B3, respectively.

Results showed that the resistance determinants varied according to the group. Each group was associated with *penA* alleles types: *penA*.2 with group A; *penA*.19 with group B1; *penA*.14 with group B2; and *penA*.22 with group B3. The *ponA* gene contained predominantly a mutation L421P in all groups except group B2, in which the majority was WT (11/18). The two isolates carrying the mutated *porB1b* (A121S) were found in group B1, and *porB1a* allele was predominant in B3 and B2. However, *porB* was predominantly WT in group A. The three isolates carrying a mutation 57Adel in *mtrR* promoter associated or not with G45D in *mtrR* coding sequence were grouped together in group A. The *mtrR* gene was WT for all isolates in B3 and isolates carrying the *mtrR* coding sequence mutation A39T were exclusively grouped into B1 and B2. The resistance determinants to CIP were quite similar in all groups except in group A where parC_S87R was predominant (6/8). The isolates carrying TEM-135 were clustered in group A (3/6) and B3 (3/6). All isolates of group B3 carried the *tetM* gene.

Groups A and B3 included only previously described STs. All ST9903 and ST7822 were grouped in groups A and B3, respectively. Group B2 included the two predominant STs, namely ST unk_1 and ST1931 while all ST 1588 were clustered in B1.

## 9. Discussion

We presented recent antimicrobial susceptibility profiles of *N. gonorrhoeae* isolates from Madagascar. This is to our best knowledge, among the first report from Madagascar of the last decennium. The CBC reported almost all *N. gonorrhoeae* isolates (98%) to be resistant to CIP. Conversely, resistance to ceftriaxone and azithromycin (only tested in 2019 and 2020) was detected among 4% and 3.1% of the tested isolates, respectively.

The almost complete resistance of *N. gonorrhoeae* against CIP did not come as a surprise and agrees with reports from other African countries (27, 28). The reduced susceptibility to CRO may have jeopardized the treatment of gonorrhoea. Unfortunately, we did not have treatment information nor treatment outcome data of the patients. The MIC of CRO decreased over the last three study years and interestingly an increase in MIC_AZT_ value was observed from 2019 to 2020. We hypothesise that this MIC_AZT_ may increase further after 2020 because of the increased use, misuse, and accessibility of AZT during and after the Covid-19 pandemic. Over the years, more than 94% of the isolates produced β-lactamase. We confirmed this phenotypic observation: in the subset of 38 PEN resistant isolates, we identified the *bla*_TEM_ gene in 36 (95%) of them.

The data was retrieved from routinely collected data and laboratory analysis of patients attending a medical analytical laboratory where biomedical analysis are paid for. In addition, the majority of the patients came from Antananarivo, the capital of Madagascar, and location of the medical laboratory. Thus, the isolates analysed in this study are not representative for the country and its population. We were also limited in the description of the epidemiology as key variables such as symptoms, clinical signs, medication, sexual orientation, travel history, and treatment outcome, are not routinely collected and thus not available for data analysis.

To get baseline data for phenotype and genotype antimicrobial resistance data of *N. gonorrhoeae*, we aimed to determine the resistance determinants of all isolates of 2020. We included in the whole genome analysis also isolates found resistant to CRO and AZT by the CBC during previous years. Unfortunately, a very small proportion of isolates obtained over the study period was available for further analysis. In addition, less than two-thirds of retrieved isolates were viable. This illustrates the difficulties routine medical laboratories, especially in low-income settings, may encounter to store and maintain fastidious microorganisms for further surveillance purposes.

We could re-test a total of six out of the 17 isolates defined to be reduced susceptible to ceftriaxone by the CBC. All six were found susceptible with very low MIC_CRO_ values. This finding underscores the need to flag and re-test isolates with certain test results such as MICs exceeding the breakpoints, especially for ceftriaxone (29). For what concerns azithromycin, two isolates considered resistant by the CBC were re-tested and found susceptible with MIC values of 0.047 and 0.032 µg/mL. Both isolates were obtained in 2020 and may therefore explain in part an unreal rise of MIC_90_ value in 2020 compared to 2019.

We did not detect any specific genomic determinant associated with CRO, AZM, SPT resistance among the sequenced isolates which is in line with the antimicrobial susceptibility profiles obtained after re-test. We observed an incremental increase of the MIC_CIP_ according to the number of mutations in *gyrA*, *parC* genes in agreement with previous reports (10).

Almost all isolates resistant to PEN, carried the *bla*_TEM_ gene encoding β-lactamase and conferring high level resistance. Three different bla_TEM_ were identified: TEM-1 was the most prevalent and is in agreement with what is usually found (9), TEM -206 has not been described previously but it is the detection of TEM-135 which is the most worrisome.TEM-135 requires only one SNP to evolve into an Extended Spectrum β-Lactamase (*ESBL*) impacting the susceptibility to all Extended-spectrum Cephalosporins (ESC) (11). Interestingly, it was already detected in one isolate from 2018 and all TEM-135 were carried by the Asian plasmid type. This observation is similar with studies from China (30), UK (31) but in contrast with a global study (32) reporting that *bla*_TEM-135_ was associated commonly with the Toronto/Rio plasmid type. Moreover, a chromosomally mediated resistance or reduced susceptibility to PEN through the accumulation of mutations was observed in all isolates. We found an equivalent high proportion of isolates carrying the *tetM* gene acquired through conjugate plasmids and associated with higher MIC_TET_ values. Although penicillin and tetracycline are since long not used for gonorrhoea treatment, the majority of the isolates contained plasmids carrying resistance genes leading to PEN and TET resistance (n=33). The high prevalence of plasmids carrying AMR genes in *N. gonorrhoeae* isolates from low- and middle-income countries (LMICs) especially in Africa was reported previously (11, 33, 34). In addition, this finding adds to the evidence that the presence and propagation of plasmids are maintained through a selective pressure and that fitness costs are attenuated in the gonococcal population.

An accumulation of chromosomal modifications in *ponA* and *mtrR* genes was detected in more than half (n=24) of the isolates, of which two carried a mutation in *porB1b* gene. The resulting overexpression of the efflux pump and structural changes in the membrane contributes to the resistance or decreased susceptibility of a wide range of antimicrobials (9, 10).

The genome-based AMR prediction of Pathogenwatch correlated mostly well with the phenotypic resistance profile. A few discrepancies were observed in the event the resistance was plasmid-mediated. All TET susceptible isolates not carrying the *tetM* gene were classified as intermediate by Pathogenwatch due to the presence of other genetic resistance determinants. Pathogenwatch classified two isolates PEN resistant because they carried a *bla*_TEM_ gene, but E-testing defined them reduced susceptible. The opposite was also observed: two PEN resistant isolates according to the E-test were classified intermediate by Pathogenwatch due to the absence of *bla*_TEM_ gene. Notwithstanding, isolates containing β-lactamase plasmids and having a MIC below the resistance threshold have been reported previously (35).

We found a wide diversity of MLST STs (23), but two-third of the identified STs were singletons. We hypothesize that the lack of clonal transmission of STs within our study population is due to the nature of our study design and small number of isolates. However, almost half of the isolates pertained to four major MLST STs: ST 9903; ST 7822; ST1931; Unk-1. Less than half of the STs were new STs and not described previously. The STs we identified and which were present in the Pathogenwatch database, were reported previously in mainly Europe (UK, Norway, Sweden, Portugal, Greece) but also in Australia, USA, and Vietnam (36). Among the most common STs in our study population, ST 7822 was prevalent in France since 2018 (37). The ST 1588 and 1931 have been described in Africa, namely in Burkina Faso, South Africa and Kenya (38–40). We hypothesize that these STs may have been imported by visitors or persons who travelled to and from Europe or the African continent. On the other hand, the observation of STUnk_1 as one of the predominant STs suggests the presence and the circulation of a possible new clone in Antananarivo, Madagascar. Fortunately, we did not identify STs such as ST9363, ST1901, associated with high-level resistance to CRO and AZT among our study population. However, we detected two CRO-susceptible isolates belonging to ST7827, a sequence type described to be associated with ESC resistance (41).

Core genome phylogenetic analysis showed that the isolates pertained to two major lineages (A and B), with one (B) presenting three clusters (B1, B2, B3). *PenA* alleles and MLST were clustered into those four groups of which *penA* 2.002 and MLST 7822 clustered in group A and penA 22.001 and MLST 9903 in B3. We did not observe a clustering of the isolates associated with their phenotypic AMR profiles. This observation may not agree with what has been reported previously (42), and is probably due to the obtained dichotomy: all isolates were fully susceptible to three antimicrobials and almost fully resistant to the other three. But we observed a clustering of some genetic resistance determinants: all *parC* S87R, *mtrR* G45D and *mtrR* promoter 57Adel were clustered only in group A, and *bla*_TEM-135_ was only found in group A and B3.

The national treatment guidelines recommend nowadays CRO eventually in combination with AZT if a co-infection with *Chlamydia trachomatis* is suspected. Although, routinely collected and reported data to GLASS may suggest that CRO is no longer a valid antibiotic for first line treatment in Madagascar (43), we could not confirm this finding. We did not detect resistant CRO among the re-tested isolates using E-test nor did we detect resistance associated mechanisms. Besides, our findings support the need to assign reference laboratories for surveillance of antimicrobial susceptibility of *N. gonorrhoeae* in order to support the ministry of health and other official bodies in updating gonorrhoea management guidelines. The benefit of re-testing by reference laboratories was already previously concluded, albeit in a high resource context, by Harris et al, as they observed that discrepancies between phenotype and genotype data were solved after re-testing of isolates (44).

We acknowledge that our study has several limitations. First of all, our epidemiological data is very limitedand does not include information required to describe the epidemiological characteristics of gonorrhoea. Secondly, we do not know whether some of the patients were related, and we lack knowledge about the patient’s symptoms, clinical signs and treatment outcome. Our study was restricted to the isolates which were remained viable after storage and may have introduced a selection bias: only more robust ones may have been included in the study.

In conclusion, efficient epidemiological or public health surveillance including data collection on gonorrhoea prevalence, incidence, and antibiotic susceptibility is required for monitoring the infection and its antimicrobial resistance profile evolution. The surveillance of *N. gonorrhoeae* is mainly laboratory-based requiring culture and antimicrobial susceptibility testing for AMR phenotyping. Consequently, *N. gonorrhoeae* surveillance is absent or not well conducted in many countries in Africa. We demonstrated with this study that genetic detection of mechanisms associated with resistance is reliable and feasible in a LMIC. But, WGS capabilities in LMICs are lacking and need to be urgently addressed. Furthermore, with appropriate epidemiological data a more comprehensive genomic epidemiological surveillance may be envisaged. However, methods for the concentration and specific capture of *N. gonorrhoeae* DNA should be developed so that WGS is fully culture independent and applicable in resource constraint settings.

## 10. Author statements

### 10.1 Author contributions

LFR, MANR, TC designed the study

ERH, FR: responsible for the routine analysis at the CBC and contributed to the data

LFR performed additional laboratory testing

LFR and MANR conducted the bioinformatic analysis

AMS preformed whole genome sequencing and made the data available to the study group

LFR and TC wrote the original draft, all contributed with review and editing

### 10.4 Ethical approval

Ethical clearance was waived by the national ethical committee of Madagascar for routinely collected and coded data analysed for surveillance purposes. Therefore, ethical clearance was not needed in this study.

### 10.5 Consent for publication

Not applicable

## Supporting information

Supplementary Tables_AMR profile NG_Mada

## 10.6 Acknowledgements

The authors wish to express their gratitude to Jean-Marc Collard, Researcher in Enteric Bacterial Pathogens Unit & French National Reference Center for Escherichia coli, Shigella and Salmonella, Institut Pasteur in Paris, France, for facilitating the contacts with NICD. We extend our thanks to Magnus Unemo, WHO centre for gonorrhoea and other Sexually Transmitted Infections, Örebro University, Örebro, Sweden, and the team of the STI reference laboratory at the Institute of Tropical Medicine, Antwerp, Belgium, for having send us the WHO and other quality control strains. A very grateful thanks to Irith De Baetselier for her critical reading of the first draft of the manuscript.

